# *Virectaria stellata* (Sabiceeae-Rubiaceae), a new sandstone cliff species from the Republic of Guinea with stellate hairs recorded for the first time in the Rubiaceae

**DOI:** 10.1101/2024.04.30.591914

**Authors:** Faya Julien Simbiano, Xander M. Van Der Burgt, Iain Darbyshire, Pepe M. Haba, Gbamon Konomou, Martin Cheek, Charlotte Couch, Sékou Magassouba

**Author notes:** Corresponding authors: M. Cheek, F. J. Simbiano.

## Abstract

*Virectaria* is a morphologically isolated genus of tropical African herbs or subshrubs, occurring from Senegal to Tanzania. *Virectaria stellata*, a new species from Guinea, is published. It is a perennial herb, with stems becoming creeping and rooting, to 60 cm long. *Virectaria stellata* has stellate hairs, recorded here for the first time in the family Rubiaceae. We hypothesize that the stellate hairs of this species result from horizontal gene flow from an Acanthaceae, most likely *Barleria*, due to their common and perhaps unique microstructure. *Virectaria stellata* is found in Forécariah and Kindia Prefectures in the Republic of Guinea. The species occurs in fissures in vertical sandstone rock at altitudes of 450 to 910 m, in sun or half-shade. An overview of sandstone endemic plant species in the vicinity of the new *Virectaria* is provided. No threats have been observed, therefore, *Virectaria stellata* is provisionally assessed here as Least Concern (LC).

## INTRODUCTION

Among the specimens collected during botanical surveys aimed at establishing Important Plant Areas in the Republic of Guinea (henceforth Guinea; Couch et al. 2019; Darbyshire et al. 2017) was a new species of *Virectaria* Bremek. (Rubiaceae: Sabiceeae) found on the Benna Plateau and Mont Kouroula in the prefectures Forécariah and Kindia respectively. The new material from Guinea was placed in *Virectaria* due to the presence of several traits that characterise this morphologically isolated genus: the stigma is unlobed and capitate and only slightly wider than the style; both style and stamens are exserted about as long as the corolla lobes; one of the two fruit valves is deciduous, the other persistent; and the floral disc is cone-like, accrescent and dehiscing into two halves in fruit; the leaves have raphides present.

*Virectaria* Bremek. (Bremekamp 1952) was erected to contain most of the African species previously referred to the genus *Virecta* L.f. (Linnaeus 1782). The International Plant Names Index (IPNI 2024) lists 28 names under *Virecta*. The neotropical *Virecta* names, together with the type of *Virecta, V. biflora* L.f., are referrable to *Sipanea* Aubl., a genus of about 19 species in northern S. America and C. America. The Asian names of *Virecta* refer to *Ophiorrhiza* L., a genus of about 320 species occurring from India to N.E. Australia and Japan. Some African *Virecta* names are referred to *Pentas* Benth., *Parapentas* Bremek (*Virecta setigera* Hiern); or *Sabicea* Aubl. (*Virecta lutea* G. Don). Verdcourt (1953) revised *Virectaria*, recognising five species from the 12 *Virecta* names attributable to the genus *Virectaria*.

*Virectaria* was often formerly place in a loosely circumscribed Hedyotideae (e.g. Hepper, 1963) together with *Oldenlandia* L., *Pentas, Parapentas* and *Hekistocarpa* Hook.f. However, Verdcourt (1953) did not concur and erected the mono-generic tribe Virectarieae (Verdcourt 1975) to accommodate the genus.

Publication and molecular placement of the monotypic Socotran *Tamridaea* Thulin & B. Bremer showed a close relationship with *Virectaria* and the placement of these two genera in an expanded Sabiceeae (Bremer & Thulin 1998). Placement of *Virectaria* within the Sabiceeae was contested by Dessein et al. (2001a) on morphological grounds, and *Virectaria* was instead placed with *Hekistocarpa* and *Tamridaea* in an expanded Virectarieae near to Sabiceeae (Dessein et al. 2001b). However, more detailed subsequent molecular studies reconfirmed placement in Sabiceeae (Khan et al. 2008a; 2008b). The monotypic Cameroonian *Hekistocarpa* is sister to the three other genera of Sabiceeae, Socotran *Tamridaea* is sister to *Virectaria*, and these two genera are in turn sister to the most species-diverse genus *Sabicea* which, with c. 167 species, extends from the Neotropics, through Africa and Madagascar, to Sri Lanka.

In this paper we describe *Virectaria stellata sp. nov*., increasing the numbers of species in the genus in Guinea from two to three (Gosline et al. 2023a; 2023b). The new species is exceptional in the Rubiaceae in having stellate hairs, and further remarkable in that they include an unusual type of stellate hair otherwise known from some Acanthaceae.

New, nationally endemic plant species continue to be steadily published from Guinea e.g. recently *Casearia septandra* <u>Breteler</u> & <u>A.Baldé</u> (<u>Breteler</u> & <u>Baldé</u> 2024, Salicaceae), *Keita deniseae* Cheek (Cheek et al. 2024, Olacaceae), *Erianthemum nimbaense* Jongkind and Phragmanthera cegeniana Jongkind (Jongkind 2023, Loranthaceae) and Gymnosiphon *fonensis* Cheek (Cheek et al. 2023, Burmanniaceae).

## MATERIALS & METHODS

All specimens cited have been seen. Herbarium material was examined with a binocular microscope fitted with an eyepiece graticule. Measurements of flower structures and fruits were made from rehydrated material. The drawings were made using the same equipment equipped with a camera lucida. Herbarium codes follow Index Herbariorum (Thiers, updated continuously). Names of species and authors follow IPNI (2024). Nomenclature follows Turland et al. (2018). The extent of occurrence and the area of occupancy were calculated using GeoCAT (Bachman et al. 2011) and the conservation assessment was made following the categories and criteria of IUCN (2012). The morphological terminology follows Beentje (2016).

## TAXONOMIC TREATMENT

Key to the species of the genus *Virectaria* in Guinea

1. Hairs stellate ………………………………………….. *V. stellata* Hairs simple ………………………………………….. 2
2. Leaves elliptic to lanceolate; hairs on stems erect, to 2 mm long …………. *V. multiflora* Leaves ovate-oblong, elliptic or sub-spathulate; hairs on stems appressed, to 0.2 mm long …………………………………………………………………………………………………… *V. procumbens*

***Virectaria stellata*** Cheek, I.Darbysh. & Simbiano **sp. nov**.

Type: Republic of Guinea, Forécariah Prefecture, Benna Plateau, 4 km West of Gombokori, 9° 44’ 19.2” N, 12° 49’ 32.9” W, 780 m, fl. & fr., 1 Nov. 2019, *Burgt & P*.*M. Haba* 2332 (holotype HNG; isotypes B, BR, EA, FI, K001381504, K001381505, LISC, MO, NY, P, PRE, SERG, SING, SL, US, WAG).

### Diagnosis

*Virectaria stellata* differs from the other species of *Virectaria* Bremek., and from all other Rubiaceae species, in the presence of stellate hairs, which are abundant on the stems, both surfaces of the leaf, and on the outer surfaces of the calyx-hypanthium and the corolla. Morphologically, *Virectaria stellata* resembles *V. tenella* J.B.Hall, a species from Ghana (Hall 1972), that is also found in vertical rock habitat. The leaf blade of *V. stellata* is 10–30 (–50) mm long(vs (5–) 7–12 mm long) and the corolla of *V. stellata* is densely stellate hairy outside (vs glabrous).

### Description

Perennial herb, prostrate, stoloniferous. Stellate hairs dense on stems, stipules, leaves (both surfaces), pedicel, calyx and corolla outer surfaces, and fruits; hairs white to colourless, 0.3–0.8 mm in diam., 10–25-armed, individual arms unequal in length, overlapping with adjacent hairs; in some stellate hairs, the central arm is a little longer to much longer than all other arms, extending from the centre of the stellate hair, 0.6–5 mm long, erect, or appressed and directed to the leaf apex (Fig. 1B). Stems drying pale reddish, initially erect, then creeping, pendant, terete, each 10–60 cm long; dense stellate indumentum; young stems herbaceous, older stems woody, 0.7–1.5 mm in diam., to 4 (–10) mm in diam. at base, internodes 0.5–4 (– 8) cm long, sometimes rooting at the nodes. Roots to 3 mm thick, to at least 50 cm long, with numerous wiry root branches, themselves highly branched. Stipules triangular, to 1 mm × 1.5 mm, bifurcate to the base into two triangular parts, apices acute to rounded; dense stellate indumentum; colleters inconspicuous. Leaves opposite, in equal pairs, decussate; petiole canaliculate, dense stellate indumentum, 2–10 × 0.5–0.7 mm, articulated at junction with stem. Leaf blades papery, drying pale green, dense stellate indumentum on both surfaces, leaf blade ovate, 10–30 (–50) × 6–17 (–28) mm, apex acute to attenuate, base obtuse to rounded; lateral nerves 4–7 on each side of the midrib, starting at 60°–70° near the petiole, starting at c. 45° towards the apex, straight, then arching upwards and becoming parallel with the margin, not uniting; nerves channelled above, raised below, domatia absent, secondary and tertiary nerves moderately conspicuous above, not branching, not reticulate, quaternary nerves not visible. Inflorescence a densely branched cyme, axillary or terminal, 1–15(–30)-flowered, dense stellate indumentum, bracts elliptic, 2–3 mm long. Flowers 13–15 × 7–9 mm. Pedicel terete, shortly cylindrical, 1–1.5 mm long, dense stellate indumentum. Calyx-hypanthium shortly cylindrical to ellipsoid, 1.2–1.5 × 0.9–1 mm, dense stellate indumentum, lacking surface sculpture; calyx tube very short or absent, calyx lobes 5, green, narrowly triangular to linear, accrescent, very slightly unequal in length, 1–2 × 0.5 mm, colleters inconspicuous, dense stellate indumentum outside, glabrous inside. Corolla white, 7–8 mm long, 8–9 mm diam. at anthesis, tube 3–5 mm long, 0.5–0.6 mm wide at base, widening gradually to 1 mm wide near lobes; lobes 5, lanceolate-oblong, 3.5–4 × 0.8–1.2 mm, apex acute, corolla outside with dense stellate indumentum, inside glabrous. Stamens 5, exserted, inserted in corollamouth, 5–6 mm long, filaments white, terete, glabrous, anthers medifixed, dark brown to black, lanceolate-elliptic, 0.9–1.1 mm long. Disc conical, purple, 0.25 × 0.1 mm, glabrous. Style white, filamentous, 9–11 mm long, exserted for half its length, glabrous. Stigma white, capitate, 0.3 mm diam.; ovary 2-locular, each locule with numerous ovules Fruit straw-coloured, overall c. 4 × 1.5–2.5 mm in diam., fruit body ellipsoid, surmounted by the accrescent calyx, lobes linear-ligulate, 0.5–1.8 × 0.2 mm, dense stellate indumentum outside, glabrous inside; fruit dehiscing longitudinally into two valves, with one valve falling, the other remaining attached to the pedicel (Fig. 1 H). Placentas 2, c. 0.7 × 0.5 mm, protruding into the locule, each bearing 20–30 seeds. Seeds bright brown, truncated-obconic, (3–) 4–6-sided, widest distally, tapering to the hilum, c. 0.5 mm × 0.2–0.25 mm, surface tuberculate, distal surface with 20–30 rounded tubercles (Fig. 1I), hilum slightly raised, orbicular, 0.025 mm diam.

**Figure 1.**
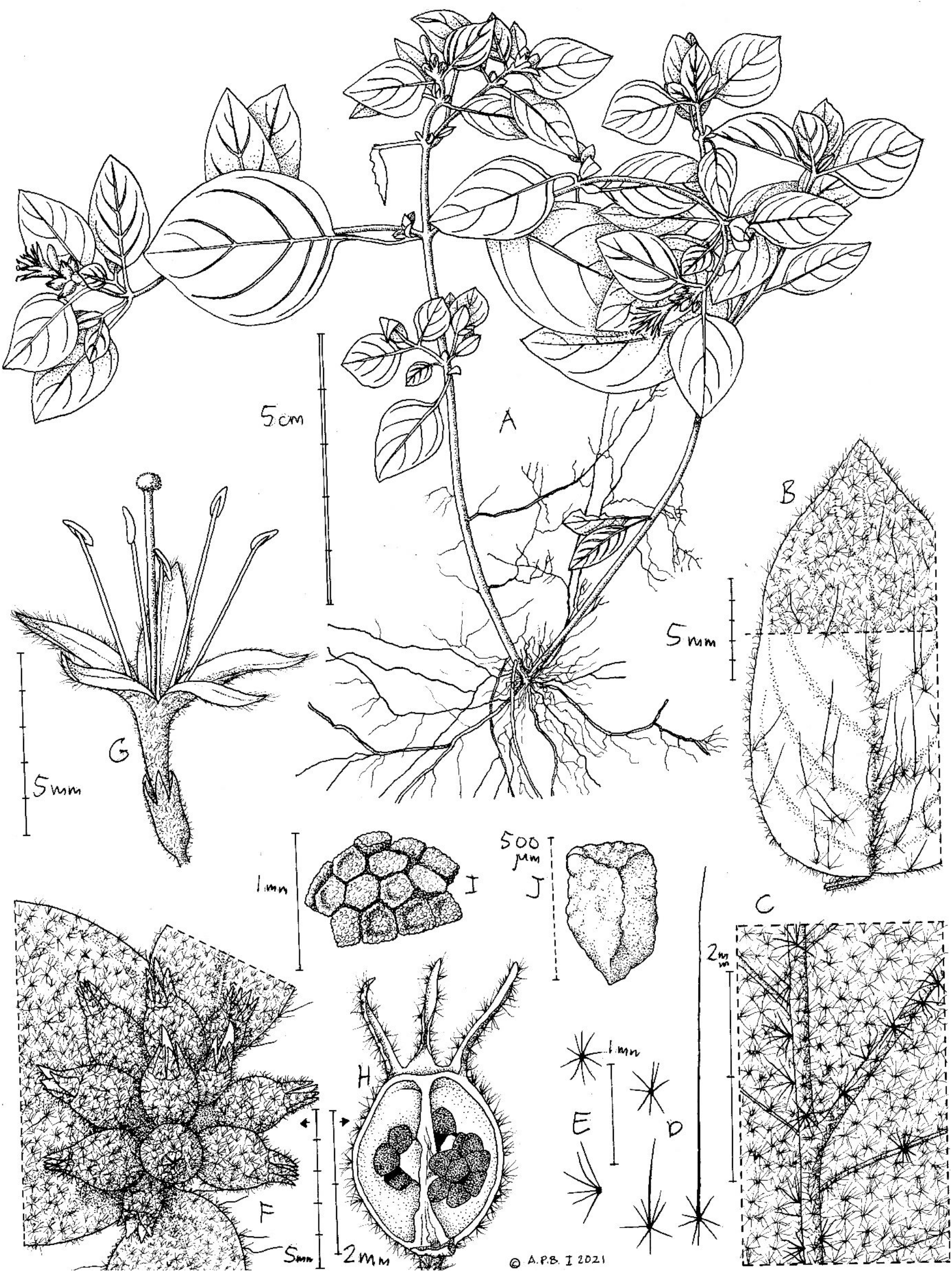
*Virectaria stellata*. A. habit, flowering plant; B. leaf upper surface (full cover of stellate hairs drawn only on distal part); C. leaf lower surface, showing midrib and secondary nerves; D. stellate hairs from upper leaf surface; E. stellate hair seen from above and side; F. infructescence, immature; G. open flower; H. fruit, after dehiscence (valve fallen); I. mature seeds *in situ* on placenta; J. mature seed, side view. Drawn from *Burgt* 2332 & 2345 by Andrew Brown.

### Etymology

The species epithet *stellata* is named after the stellate hairs that are so characteristic of this species (Figs. 1&3).

### Distribution

Endemic to Republic of Guinea, Kindia and Forécariah Prefectures (Map 1).

### Habitat & ecology

*Virectaria stellata* occurs in fissures on vertical sandstone rock at altitudes of 450 to 910 m, in sun or half-shade (Fig. 2A). The species has been found growing with other Guinea endemic species, *Cailliella praerupticola* Jacq.-Fél. (Melastomataceae) (*Burgt* 2332), *Fleurydora*

**Figure 2.**
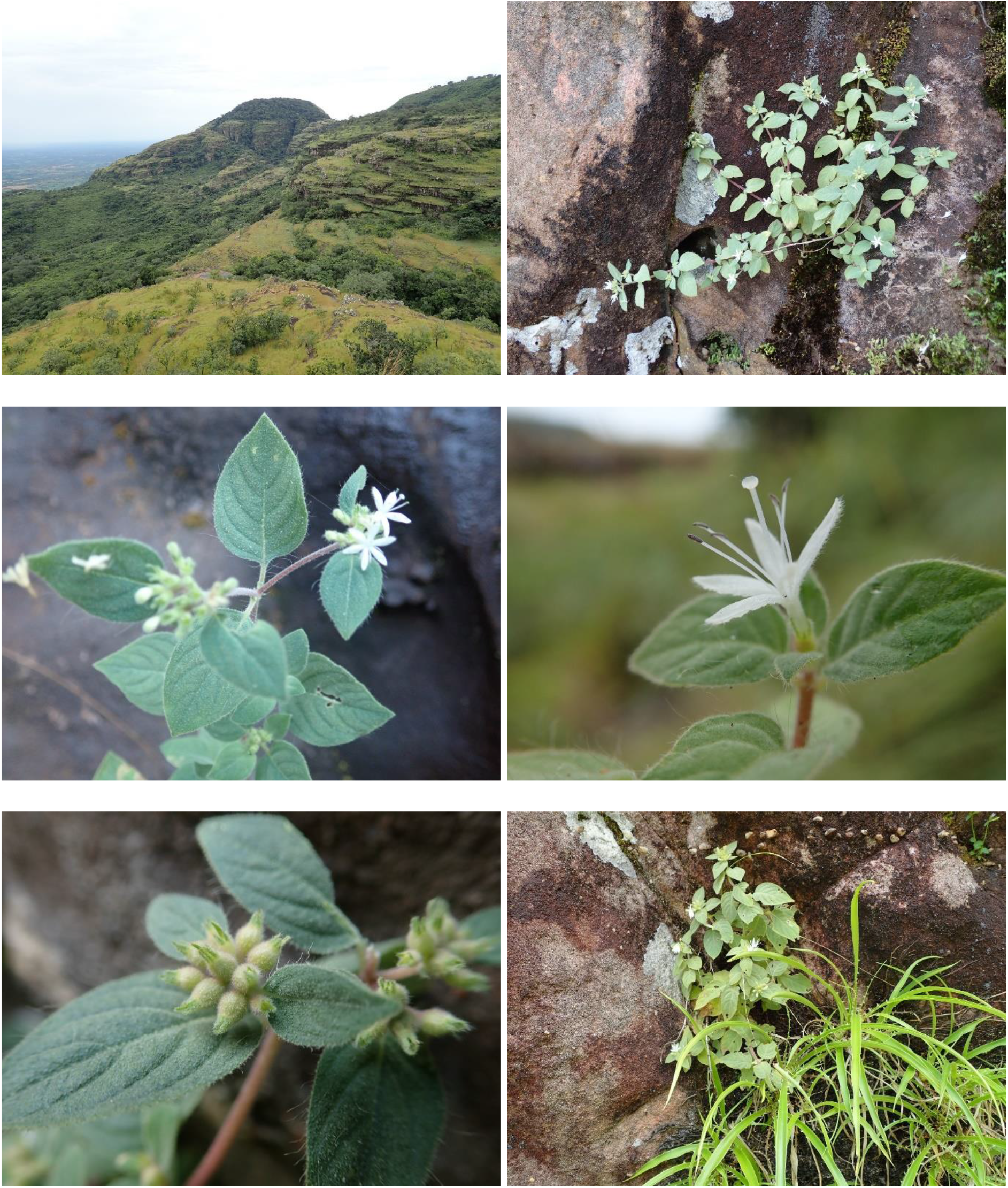
*Virectaria stellata*. A. Vertical rock habitat on the Benna Plateau. The type was collected on the rocks at the centre of the photo; B. Flowering plant; C. Inflorescence; D. Flower; E. Young fruits; F. Flowering plant growing with *Pitcairnia feliciana*. B, D, E, F from: *Burgt* 2332; C from *Simbiano* 683. Photos: A, B, D–F Xander van der Burgt; C Faya Julien Simbiano.

*felicis* A.Chev. (Ochnaceae) (*Burgt* 2332), *Kindia gangan* Cheek (Rubiaceae) (*Burgt* 2345) and *Pitcairnia feliciana* (A.Chev.) Harms & Mildbr. (Bromeliaceae) (*Burgt* 2332; Fig. 2F).

### Phenology

*Virectaria stellata* was collected in flower in November, and in fruit in February, March and November. The dry season is from November to March and the rainy season is from April to October.

### Conservation status

*Virectaria stellata* is known from eight collections in three localities which correspond to three locations. Three collections are from the centre of the Benna Plateau, South of Kindia, where thousands of plants were seen. Three other collections are from the 250 m high vertical rock escarpment at the Northern end of the Benna Plateau, South of Kindia; where hundreds of plants were seen. Many more plants are probably present here, higher up on the vertical rocks, where they cannot be seen from the base of the vertical rock. Two collections are from Mont Kouroula, North of Kindia, about 50 km North of the Benna Plateau; thousands of plants were seen here. The extent of occurrence (EOO) is 121 km^2^, and area of occupancy (AOO) is 20 km^2^. Although suitable areas of sandstone cliff habitat exist where the species is not found, it is likely that the EOO and AOO underestimate the true distribution of the species, as it is likely that other populations exist but have not yet been found. The extent of occurrence of the species is supposed to probably be less than 5,000 km^2^ and the area of occurrence less than 500 km^2^. Fires started by herders during the dry season have been observed to damage plants at the base of the cliffs on which this species grows. Fires are also set up cliffs to drive off bees from their nests, to collect honey (Cheek pers. obs., Kindia region). Although underground perennating structures are not recorded in this species, burned plants were observed by the specimen collectors to resprout from the base. Some plants have buds at the stem base showing signs of regeneration after the passage of fire (see for example *Burgt* 2332, HNG, K, SERG). Juvenile plants and young flowering plants were recorded near the base of the cliffs, in areas presumably affected by dry-season fires. It seems that, when mature plants are destroyed by fire, a new generation will grow back in the same place.

Therefore, no decline was observed in area, extent and/or quality of habitat, as well as in number of mature individuals. *Virectaria stellata* is therefore provisionally assessed here as Least Concern (LC).

### Additional specimens examined

**GUINEA. Forécariah Prefecture**, Benna Plateau, 5 km W of Gombokori, 9° 44’ 22.9” N, 12° 50’ 11.4” W, 910 m, fl. & fr., 4 Nov. 2019, *P*.*M. Haba & Burgt* 1363 (B, BR, G, HNG, K, MO, P, SERG, US, WAG). **Kindia Prefecture**. Mont Kouroula, 6 km E of Koumbaya, 10° 10’ 57” N, 12° 49’ 21.4” W, 450 m, fl. & fr., 7 Nov. 2019, *Burgt, P*.*M. Haba & Holt* 2345 (HNG, K, P, WAG); Mont Kouroula, 6 km E of Koumbaya, 10° 10’ 52.5” N, 12° 50’ 06.5” W, 460 m, fr., 10 Nov. 2019, *P*.*M. Haba, Burgt & Holt* 1375 (HNG, K); near Mambia village, mount Yon-Ya, 9° 46’ 31.1” N, 12° 48’ 8.9” W, 720 m, fr., 30 March 2023, *Konomou, Burgt, Conté & Thiam* 1128 (B, BR, G, HNG, K, MO, P, PRE, SERG, WAG); near Molota, path to Balaqui, 9° 46’ 23.6” N, 12° 49’ 57.6” W, 660 m, sterile, 31 March 2023, *Thiam, Konomou, Conté & Burgt* 23 (HNG, K); between villages Dokokouré and Tinekouré, 9° 46’ 43.6” N, 12° 49’ 34” W, 640 m, fl., 16 Nov. 2023, *Simbiano, Thiam, Touré & Bangoura* 683 (HNG, K).

**Map 1.**
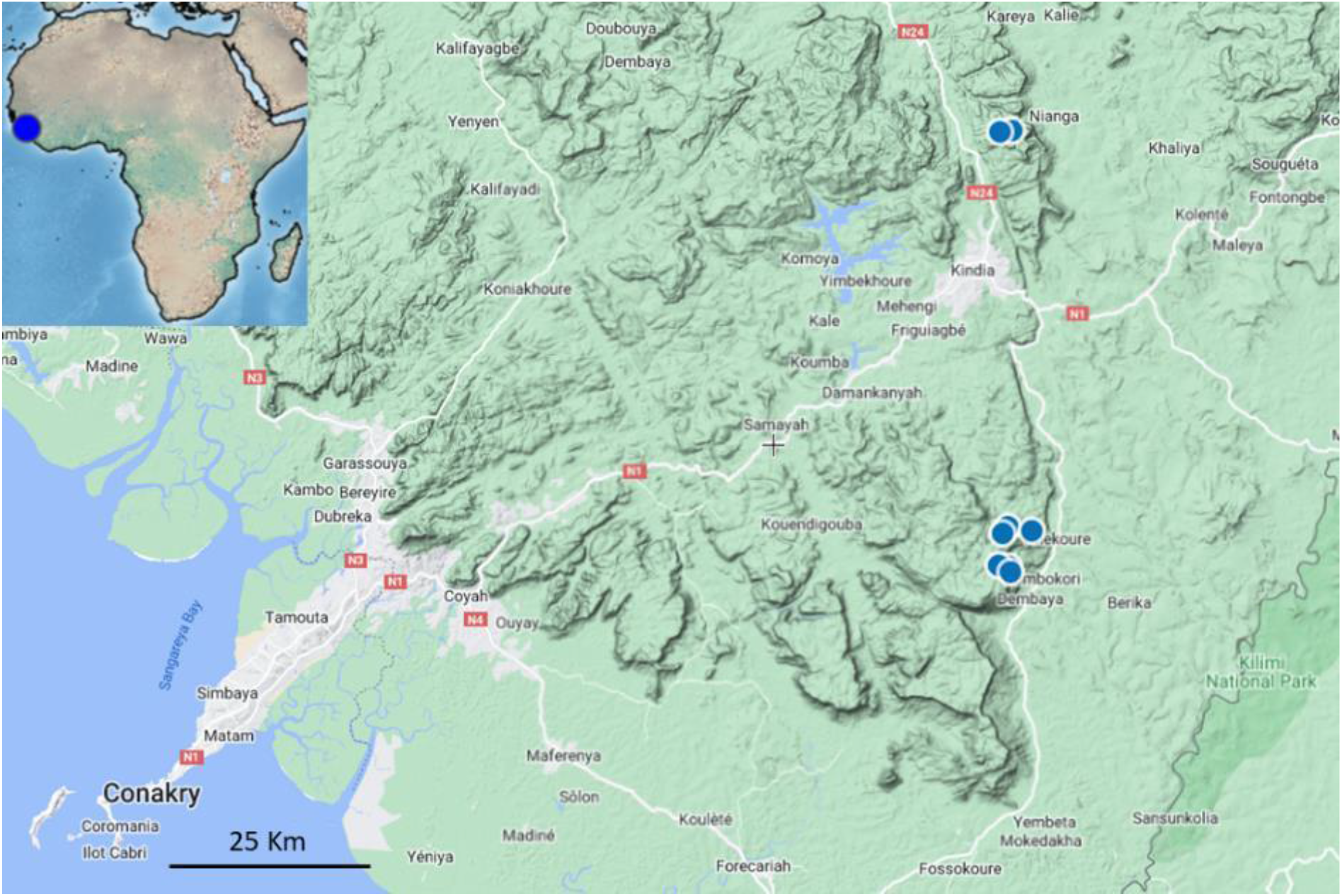
Distribution of *Virectaria stellata* (blue dots). Map data © Google 2024.

## DISCUSSION

### Stellate hairs: a newly discovered trait for Rubiaceae

The presence of stellate hairs in *Virectaria stellata* (Fig. 1 and 3) is remarkable because not only is it here recorded for the first time in the genus *Virectaria*, but also in the entire family Rubiaceae, from which, until now, only simple hairs have been recorded (Robbrecht 1988). In some of the stellate hairs, the central arm greatly exceeds the others, by a factor of ten or more. On the leaf upper surface, these stellate hairs are appressed and directed towards the leaf apex. This unusual stellate hair type is otherwise known to us from the tribe Barlerieae in Acanthaceae, in the genera *Barleria* L. and *Lepidagathis* Willd. (Darbyshire et al. 2010). In *Barleria* in particular, this trait has evolved independently in several lineages (Balkwill & Balkwill 1997; Darbyshire et al. 2019; Comito et al. 2022). Two species of *Barleria* in Guinea have stellate hairs, *B. asterotricha* Benoist and *B. maclaudii* Benoist, and those of *B. asterotricha* in particular are similar to the hairs seen in *Virectaria stellata*. We conjecture that horizontal gene transfer from a *Barleria* to the progenitor of *V. stellata* may be the explanation for this phenomenon, in the absence of any other explanation. We speculate that genetic material may have been transferred by a sap-sucking insect moving from feeding on a *Barleria* to a *Virectaria*.

**Figure 3.**
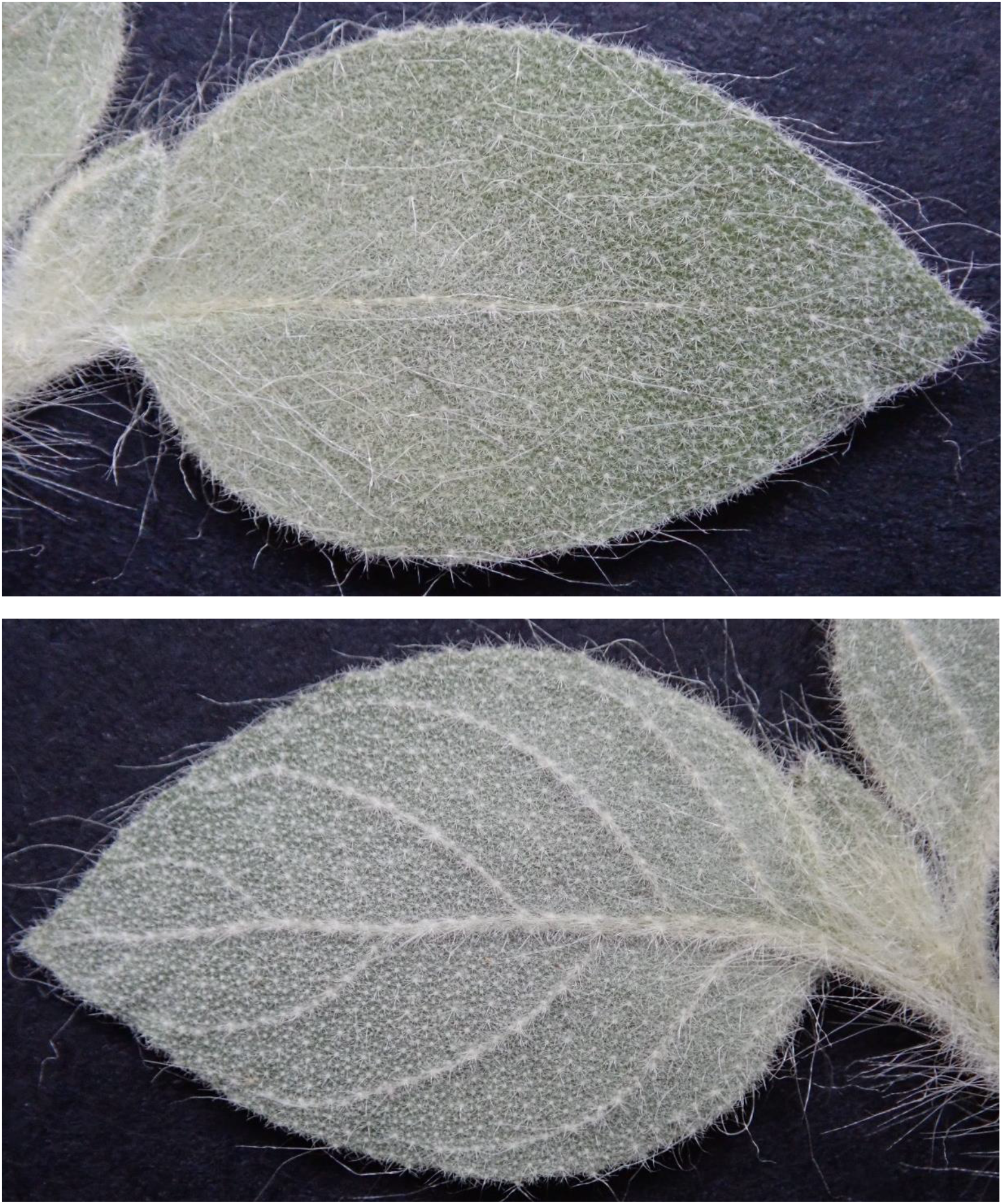
*Virectaria stellata*. A. Adaxial side of the leaf with stellate hairs, including many with one arm to 5 mm long; B. Abaxial side of the leaf with stellate hairs, including a few with one arm to 5 mm long. Length of leaf blade 19 mm. From *P*.*M. Haba* 1375. Photos: Xander van der Burgt.

### Endemics of the sandstone table mountains of the Fouta Djalon

Of the 22 Important Plant Areas (IPAs) in Guinea (Couch et al. 2019), five are located in the sandstone table mountains area. Although *Virectaria stellata* does not occur in any of these five IPA’s, one of the three localities where the species is found, is on Mont Kouroula, located in the buffer zone of the Mont Gangan IPA area (Couch et al., 2019). The Benna Plateau contains two of the three localities where *Virectaria stellata* occurs, is rich in rare plant species, and should be considered for designation as an IPA. For further information on the sandstone habitats of the Fouta Djalon see Couch et al. (2019: 20–29).

*Virectaria stellata* is the latest in a steady flow of new species to science published from the sandstone habitats of the southwestern outliers of the Fouta Djalon highlands in Guinea. The table mountains are perhaps best known to botanists for being the home of *Pitcairnia feliciana* (Bromeliaceae), the only Old World species of that family (Larridon 2018). *Fleurydora felicis* (Ochnaceae) a monotypic tree genus arising from a South American clade, has a similar geographic range and is also restricted to sandstone cliffs (Canteiro & Cheek 2019). Many other endemic plant species occur on the table mountains. These include the monotypic genus *Benna alternifolia* Burgt (Melastomataceae), *Cailliella praerupticola* (Melastomataceae), *Ctenium bennae* Xanthos (Poaceae, Xanthos et al. 2021), *Gladiolus mariae* Burgt (Iridaceae), *Impatiens bennae* Jacq.-Fél. (Balsaminaceae), *Inversodicrea tassing* Cheek (Podostemaceae, Cheek et al. 2019), *Kindia gangan* (Rubiaceae, Cheek et al. 2018), *Mesanthemum bennae* Jacq.-Fél. (Eriocaulaceae), *Rhytachne perfecta* Jacq.-Fél. (Poaceae), *Tephrosia kindiana* Haba, B.J.Holt & Burgt, *Ternstroemia guineensis* Cheek (Pentaphylacacae, Cheek et al. 2020), and *Trichanthecium tenerium* Xanthos (Poaceae, Xanthos et al. 2020). There is no doubt that more discoveries of new taxa, and range extensions, will be made if botanical survey work continues.

## ACKNOWLEDGEMENTS

Five collections of *Virectaria stellata*, including the type collection, were made during two seed collection expeditions for the Global Tree Seed Bank project of the Millennium Seed Bank Partnership of Kew, funded by the Garfield Weston Foundation. Three collections were made during plant conservation work funded by the Fondation Franklinia project “Conservation of threatened trees species in three Tropical Important Plants Areas of Guinea”, and the Darwin Initiative of the Department of the Environment Food and Rural Affairs (DEFRA), UK government (project Ref. 23–002). Mr Abdoulaye Yéro Baldé, former Minister, Guinean Ministry of Higher Education and Scientific Research, Dr Binko Mamady Touré, former Secretary General of the same Ministry, and Dr. Facinet Conté, Secretary General of the same Ministry, are thanked for their cooperation. Colonel Layaly Camara, former Director, Direction National des Eaux et Forêts, Mr Mamadou Bella Diallo, Nana Koulibaly, T. Delphine Kolié, and Mr Alpha Illias Diallo, CITES Focal Point, Direction National des Eaux et Forêts, authorised the export of the plant specimens. The first author’s training visit to Kew to write this paper was funded by the JRS Biodiversity grant (70022) “Enhancing data access to transform Guinea’s capacity to identify and protect its threatened plants”. The Prefects of Forécariah and Kindia Prefectures are thanked for their hospitality during the fieldwork. Two anonymous reviewers are thanked for constructive comments on an earlier draft of the paper.

## Declarations

The authors declare that they have no conflict of interest.

